# A kinematic analysis of *Micrurus* coral snakes reveals unexpected variation in stereotyped anti-predator displays within a mimicry system

**DOI:** 10.1101/754994

**Authors:** Talia Y. Moore, Shannon M. Danforth, Joanna G. Larson, Alison R. Davis Rabosky

**Affiliations:** Robotics Institute, University of Michigan, 2350 Hayward St, Ann Arbor, MI 48109; Ecology and Evolutionary Biology, University of Michigan, 1105 N. University Ave, Ann Arbor, MI 48109; Museum of Zoology, University of Michigan, 3600 Varsity Drive, Ann Arbor, MI 48108; Mechanical Engineering, University of Michigan, 2350 Hayward St, Ann Arbor, MI 48109

**Keywords:** aposematism, biomechanics, coral snake mimicry, curvature, Elapidae, non-locomotory motion, Peruvian Amazon, snake behaviour

## Abstract

1. Warning signals in chemically defended organisms are critical components of predator-prey interactions, often requiring multiple coordinated display components for a signal to be effective. When threatened by a predator, venomous coral snakes (genus *Micrurus*) display a vigorous, non-locomotory thrashing behaviour that has been only qualitatively described. Given the high-contrast and often colourful banding patterns of these snakes, this thrashing display is hypothesized to be a key component of a complex aposematic signal under strong stabilizing selection across species in a mimicry system.
2. By experimentally testing snake response across simulated predator cues, we analysed variation in the presence and expression of a thrashing display across five species of South American coral snakes.
3. Although the major features of the thrash display were conserved across species, we found significant variation in the propensity to perform a display at all, the duration of thrashing, and the curvature of snake bodies that was mediated by predator cue type, snake body size, and species identity. We also found an interaction between curve magnitude and body location that clearly shows which parts of the display vary most across individuals and species.
4. Our results suggest that contrary to the assumption in the literature that all species and individuals perform the same display, a high degree of variation persists in thrashing behaviour exhibited by *Micrurus* coral snakes despite presumably strong selection to converge on a common signal. This quantitative behavioural characterization presents a new framework for analysing the non-locomotory motions displayed by snakes in a broader ecological context, especially for signalling systems with complex interaction across multiple modalities.

## Introduction

Venomous prey animals often use conspicuous phenotypes to communicate their lethal toxicity to potential predators (Ruxton 2018). These aposematic signals can be visual, chemical, acoustic, or can involve complex interactions between multiple distinct components, such as colour patterning and body motion (Rowe and Halpin 2013; Dalzeill and Welbergen). According to the theory of mimicry, toxic prey animals can reinforce the aposematic signal to their predators by converging on a common phenotype, even across multimodal components (*e*.*g*., conspicuous colour and behavioural display; Müller 1878; Wallace 1867; Sherratt 2008).

Coral snakes are highly venomous elapid snakes in the genus *Micrurus* that have been described as a mimicry system using visual warning signals of their chemical defence (Campbell and Lamar 2004). Most coral snake species are found in the Neotropics, with the highest sympatric species richness in the Western Amazon Basin (Davis Rabosky et al. 2016a). Coral snakes are well known for their conspicuous red and black coloration (Fig. 1A), a high-contrast banding pattern that effectively creates an aposematic signal deterring avian (Smith 1975; Smith 1977) and potentially mammalian predators (Green and McDiarmid 1981; Savage and Slowinski 1992; Martins 1996; Buasso et al. 2006). In addition to their bright coloration, coral snakes encountering a threat also produce a distinctive anti-predator display that includes elements such as body flattening, intermittent thrashing, head hiding, and coiling of the tail, which is often elevated and waved or “waggled” (Greene 1973; Greene 1979; Fig. 1A; Supp. Video 1). In some species, this display is also accompanied by an auditory cloacal “popping” sound, eversion of the hemipenes, and emission of cloacal musk and faeces (Greene 1973; Sazima and Augusto 1991). Because such similar elements of this behavioural display have been reported across many coral snake species in both Asia and the Americas (Brown et al. 2013), this anti-predator response is expected to have 1) an old origin that predates the arrival of this clade in the Western Hemisphere ∼35mya and 2) a significant genetic basis like other homologous traits derived from shared ancestry (Wake et al. 2011). The main purpose of this display has been hypothesized as inducing a surprisingly effective cognitive illusion that reduces the ability of the attacking predator to identify and target the head (Roze, 1996). The intermittent, self-occluding thrashing is described as the foundational display component driving this effect in all species examined.

**Figure 1:**
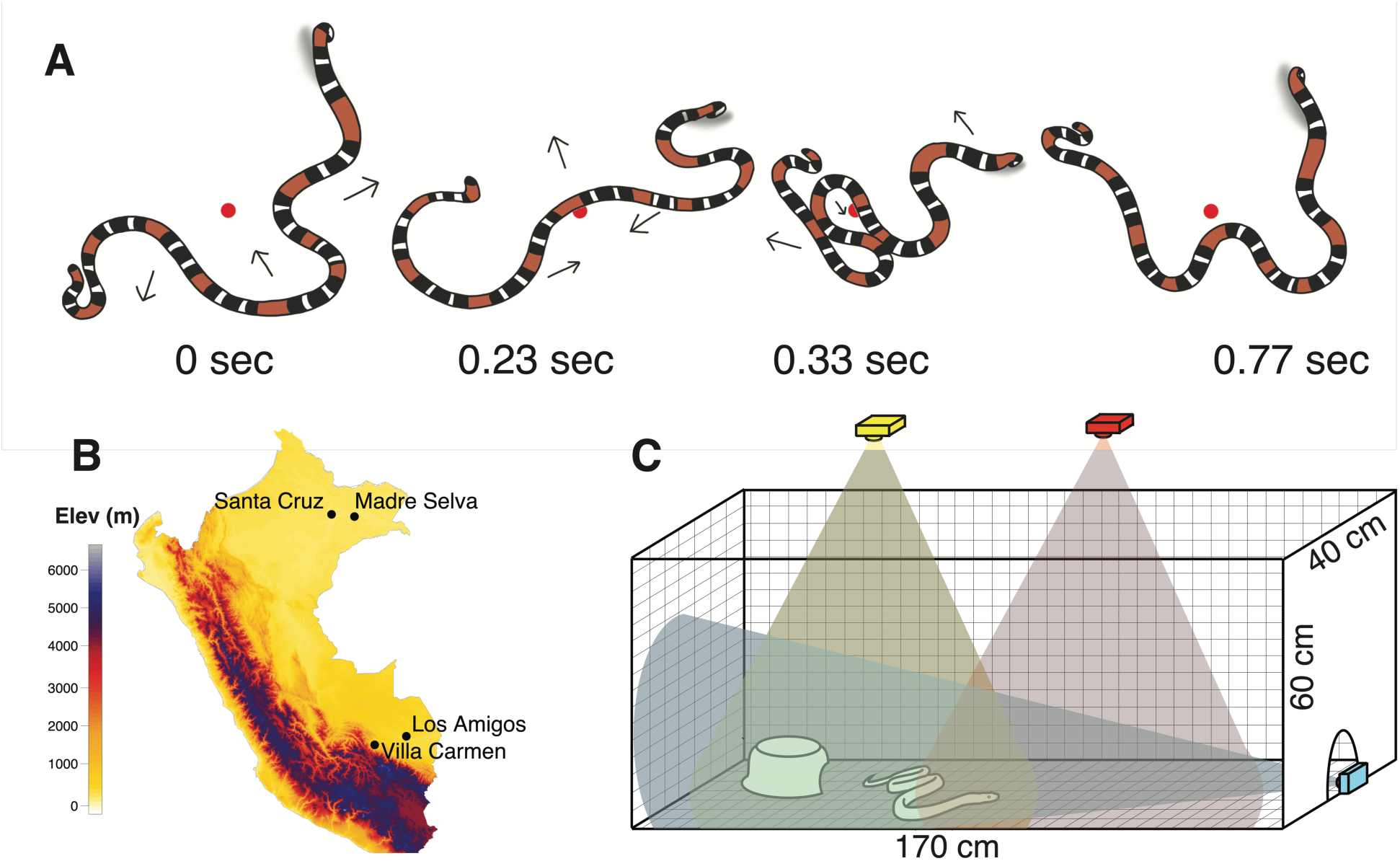
(A) An illustration of a coral snake thrashing display. Each of the four illustrations depict an individual coral snake progressing through different phases of a thrashing display (based on data in this study). The snake position with respect to the red dot shows that there is negligible translocation of the snake, despite vigorous movements. The snake head is on the left side of each illustration, and the coiled tail is elevated throughout the display. (B) A map of Peru showing our four field sites, all in lowland Amazonian rainforest. (C) A schematic of the pop-up behavioural arena. The bottom and sides of the arena are constructed from corrugated plastic and connected using brass fasteners. Lines are drawn 2cm apart using permanent ink and covered with a clear adhesive plastic (Con-Tact). Three Go-Pro cameras were positioned so that their fields of view partially overlapped and included the entire floor-space of the arena. An opaque plastic bowl with a portion removed was placed in one end of the arena to provide a refuge for the snake.

However, individual-level variation in mimicry traits is also expected to exist, with critical impacts on the ecological and evolutionary dynamics of mimicry over space and time. Variation in the banding elements of coral snake colour pattern has been well documented (Davis Rabosky et al 2016a, b) and experimentally tested using clay replicas for its effect on predator deterrence (Brodie III 1993, Buasso et al. 2006, Kikuchi and Pfennig 2010). As previous authors have noted, clay replica studies may not accurately reflect the true deterrence value of a given phenotype because they exclude behaviour (Brodie III 1993). Behavioural movement has the potential to greatly alter the perception of a colour pattern (Titcomb et al. 2013), suggesting that colour pattern and non-locomotory behaviour in coral snakes may interact to produce a complex anti-predator signal that is not fully explained by either individual component. However, neither the drivers of variation in *Micrurus* anti-predator displays nor the relationship of behaviour to colour pattern has ever been tested.

While snake locomotion has been quantitatively characterized in a variety of contexts (Lissmann 1950; Jayne 1986; Moon and Gans 1998; Hu et al. 2009; Socha 2011, Titcomb et al. 2013), most studies of non-locomotory snake behaviours rely upon qualitative descriptions (Arnold and Bennett 1984, Brodie III 1992). Here we present the first quantitative kinematic analysis of non-locomotory anti-predator behaviours in *Micrurus* coral snakes and test for signal-reinforcing similarity within and among species, as predicted by mimicry theory. By characterizing these behaviours in response to experimentally induced predator contexts, we form a functional basis for understanding both the signalling mechanism of the aposematic phenotype and the selective pressures shaping behavioural convergence among species in a mimicry system.

## Methods

### Data collection

All animal-related procedures have been approved by the University of Michigan Institutional Animal Care and Use Committee (Protocols #PRO00006234 and #PRO00008306) and the Peruvian government SERFOR (Servicio Nacional Forestal y de Fauna Silvestre; permit numbers: 029-2016-SERFOR-DGGSPFFS, 405-2016-SERFOR-DGGSPFFS, 116-2017-SERFOR-DGGSPFFS). We collected data during five field expeditions in the Amazonian lowlands of Peru from March 2016 to December 2018, at Villa Carmen, Los Amigos, Madre Selva, and Santa Cruz Biological Stations (Fig. 1B). We captured snakes either in funnel traps or opportunistically during transects, then transported the snakes in fabric bags secured within 20L lidded buckets back to the station. During capture and handling, all trained personnel were equipped with snake hooks, tongs, venom defender gloves (1-2-1 Products Ltd., Alfreton, UK), and knee-high rubber boots to avoid envenomation.

We recorded anti-predator behaviour in a pop-up behavioural arena constructed of corrugated plastic (Fig. 1C) illuminated by a string of led lights attached to the inner surface at the top edge of the arena walls (see Davis Rabosky, Moore, et al. *submitted* for more details on construction). We marked the inner surface of the arena with visual fiducial markings to aid in the removal of lens distortion and measurement. We covered the inner surface of the arena with an adhesive transparent plastic film (Con-Tact, Rubbermaid) to facilitate thorough and rapid cleansing and preserve the visual fiducial markings. Since previous research has shown that snakes are physiologically affected by temperature, chemical cues, and light in an environment (Schieffelin and de Quieroz 1991), it is likely that the behaviours exhibited in laboratory environments and by captive individuals differ significantly from those exhibited under natural conditions. Therefore, we made every effort to ensure similar experimental conditions for each behavioural trial. After capture, snakes were kept undisturbed in bags for less than 24 hours before behavioural trials and the inner surface of the arena was washed with unscented soap and water to limit exposure to the chemical cues of previous experimental subjects.

We placed snakes into the arena, one at a time, and immediately began recording their behaviour. After the first trial, we measured surface temperature of each individual with a Raytek Raynger ST81 infrared temperature sensor. We recorded snake behaviour using either two or three GoPro (San Mateo, California) Hero 4+ Black or three Hero 5+ Black cameras filming from overhead and lateral views (see Fig. 1C for camera positions) at 30, 60, or 120 frames per second, depending on the lighting conditions.

We used three different stimuli to elicit anti-predator behaviours: overhead looming, pulsed vibration, and a tactile stimulus through physical contact. To simulate avian predation threat, we quickly moved a piece of cloth across the top of the arena to create the visual looming and pressure wave stimuli produced by a swooping bird. To simulate a large mammal predator, we used the vibration produced by a cellular phone and placed it in contact with the arena. To test for response after contact with a predator, we used a 1m snake hook to lightly tap the snake. We randomized all treatments across individuals, and we recorded snake behaviours for up to two minutes, allowing one minute of time to rest between the trials.

After behavioural testing, we either vouchered snakes into the University of Michigan Museum of Zoology (UMMZ) or the Museo de Historia Natural in Lima, Peru (MUSM), or released the snake at the point of capture. All vouchered snakes were also weighed for body mass, measured for snout-vent length (SVL) and tail length, and sexed where possible. Field numbers and museum accession numbers (when available) for each individual are reported in Table 1.

**Table 1:** Morphological and geographical information for the snakes examined in this study. Note that the Snout-Vent Length (SVL) for *Micrurus lemniscatus* Released9 was estimated from a still video frame. As snake morphometrics are taken at the time of vouchering, SVL, mass, and sex were not directly measured for released individuals.

| Species | Catalog Number | Field Number | Sex | SVL (mm) | Mass (g) | Locality | Thrashing? |
| --- | --- | --- | --- | --- | --- | --- | --- |
| *Micrurus annellatus* | MUSM 39056 | RAB2815 | F | 410 | 17.5 | Los Amigos | No |
| *Micrurus annellatus* | UMMZ 246856 | RAB1144 | F | 422 | 12.0 | Los Amigos | Yes |
| *Micrurus annellatus* | UMMZ 248451 | RAB3275 | F | 497 | 18.1 | Los Amigos | Yes |
| *Micrurus hemprichii* | UMMZ 246857 | RAB1810 | M | 740 | 86.0 | Madre Selva | Yes |
| *Micrurus hemprichii* | MUSM 37347 | RAB2035 | M | 617 | 52.5 | Madre Selva | Yes |
| *Micrurus lemniscatus* | NA | Released5 | NA | NA | NA | Villa Carmen | No |
| *Micrurus lemniscatus* | NA | Released9 | NA | 848* | NA | Villa Carmen | Yes |
| *Micrurus lemniscatus* | NA | Released12 | NA | NA | NA | Villa Carmen | Not analyzed (out of frame) |
| *Micrurus lemniscatus* | UMMZ 246858 | RAB1993 | F | 632 | 23.4 | Madre Selva | Yes |
| *Micrurus lemniscatus* | MUSM 37348 | RAB2415 | F | 372 | 6.1 | Santa Cruz | Yes |
| *Micrurus lemniscatus* | MUSM 39057 | RAB2706 | M | 816 | 68.9 | Los Amigos | Not analyzed (snake hook interference) |
| *Micrurus lemniscatus* | UMMZ 248456 | RAB2915 | F | 688 | 34.6 | Los Amigos | Yes |
| *Micrurus lemniscatus* | UMMZ 248457 | RAB3333 | M | 595 | 35.0 | Los Amigos | Yes |
| *Micrurus lemniscatus* | UMMZ 248452 | RAB3487 | F | 632 | 28.4 | Los Amigos | Yes |
| *Micrurus lemniscatus* | MUSM 39853 | RAB3573 | M | 570 | 25.7 | Los Amigos | Yes |
| *Micrurus lemniscatus* | MUSM 39854 | RAB3574 | M | 590 | 25.5 | Los Amigos | Yes |
| *Micrurus lemniscatus* | MUSM 39855 | RAB3578 | M | 453 | 14.1 | Los Amigos | Yes |
| *Micrurus obscurus* | UMMZ 246859 | RAB0665 | F | 261 | 5.2 | Villa Carmen | Yes |
| *Micrurus obscurus* | MUSM 37350 | RAB0698 | F | 237 | 5.1 | Villa Carmen | Yes |
| *Micrurus obscurus* | UMMZ 246860 | RAB1054 | M | 775 | 81.0 | Los Amigos | Yes |
| *Micrurus obscurus* | UMMZ 248458 | RAB3570 | F | 766 | 86.0 | Los Amigos | Yes |
| *Micrurus surinamensis* | UMMZ 246861 | RAB1099 | F | 657 | 100.0 | Los Amigos | No |
| *Micrurus surinamensis* | MUSM 37352 | RAB1100 | F | 391 | 25.2 | Los Amigos | Yes |
| *Micrurus surinamensis* | MUSM 37353 | RAB1101 | M | 421 | 32.5 | Los Amigos | Yes |
| *Micrurus surinamensis* | UMMZ 246862 | RAB1511 | F | 948 | 240.0 | Madre Selva | No |

### Video analysis

We selected videos for analysis that included thrashing behaviour with minimal translocation that stayed within the field of view of one camera (Fig. 1A). We used the Adobe Premiere Pro (Adobe Systems, San Jose) built-in filter for GoPro Hero 4+ Wide angle to remove lens distortion. We wrote custom Matlab (Mathworks, Natick, Massachusetts) code to perform a projective transformation on each video frame, removing perspective to produce an image for direct measurement.

We used QuicktimePro 7 to watch the videos frame-by-frame and recorded the first and last frames that included motion as the start and end of each thrash. We used ImageJ to measure the length of the snake in the video image, which was compared to measurements taken at the time of vouchering. We traced the centreline of the snake body at the end of each bout of thrashing with ImageJ (Fig. 2A).

**Figure 2:**
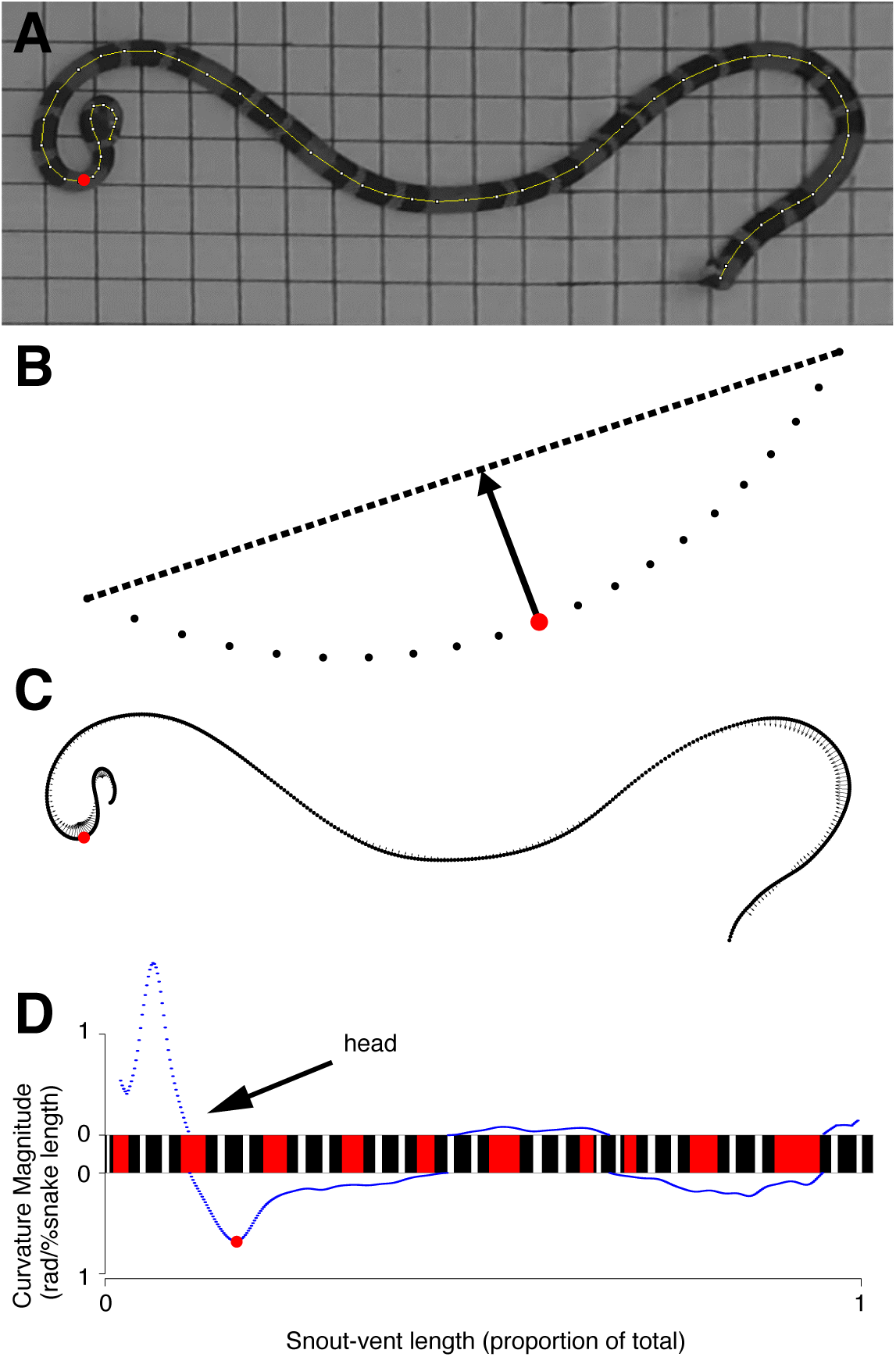
Methods for calculating the curvature of a snake. (A) Undistorted and rectified video frames at the conclusion of each bout of thrashing were traced by hand in ImageJ. (B) Curvature vectors are shown along each sampled point of the snake centreline as arrows. (C) For each focal point in the snake centreline (the red focal point in B corresponds to the red focal point in B, the curvature was calculated as the distance from the point to a line connecting the tenth points on the left and right. (D) The corresponding curvature magnitudes plotted to the right and left of a straightened depiction of a coral snake. The colour pattern of the coral snake reflects the observed colour pattern of this individual.

We wrote custom Matlab code to analyse the centreline of the snake body. First, we resampled the traced centreline to produce 500 evenly spaced points (Fig. 2B). We corrected for noise in centreline tracing by smoothing the centreline using a moving average method with a span of 30 points. Each point along this smoothed centreline can be thought of as a point on the edge of a circle that provides the closest approximation to the body curvature at that point. The angle of the tangent line of this circle may differ drastically from point to point in acutely curved areas. To calculate the curvature at each point along the body, we measured the distance from the focal point to a line formed by the tenth points to the left and right (Fig. 2C). The curvature, commonly reported in radians per unit length, is a measurement of the tangent direction’s sensitivity among nearby points on a curve. Higher curvature values indicate more acute curves.

We did not compute the curvature for the anterior and posterior extrema with fewer than ten points on a side (2% of body length). We used built-in Matlab functions to locate the areas of local maximum curvature along the body of the snake (for example, the red dot in all panels of Fig. 2). If multiple local peaks were recorded in a 30-point window, the curvatures and their indices were averaged to one point.

To automatically determine whether a particular curvature vector was located to the right or left of a snake’s head, we wrote custom Matlab code to record the angle of each curvature vector in a snake-centreline coordinate frame. We defined the temporary snake-centreline axis as a vector pointing from the current point of interest towards its neighbouring point, in the direction closer to the head of the snake. A curvature vector falling to the left of the snake-centreline axis indicated a curve to the left of the snake’s head (Fig. 2, red dot). With this method for computing curve direction, we calculated the percentage of left or right instances of local maximum curvature in each observation.

### Statistical approach

To maximize our inference ability and include information from all individuals, we first assessed whether snakes displayed any anti-predator behaviour (e.g. thrashing, escaping, or head hiding) in every recorded trial of *Micrurus* behaviour as a binary variable (presence or absence of a response). For the individuals that did respond, we then assessed the expression of that response using the number of thrashing events, their durations, and the body location and direction of the curvatures in each post-thrash pose. For the purpose of statistical modelling, we calculated the sum of the magnitudes of curvature for each of the 500 points along the body of the snake for each post-thrash pose, and we quantified each individual’s preference for a left or right head kink by determining the direction of the most anterior curve across multiple observations in the same trial. For each of these response variables, we constructed generalized linear mixed models (GLMMs) to test for effect of species, body size (SVL), sex, and stimulus type on the presence or expression of a response while accounting for individual collection ID as a random effect because every individual was tested more than once. For binary (Y/N) response tests of stimulus, we only included the treatments that had more than 5 observations per response category (contact, looming, and vibration), which removed 12 trials of 160 total. We also tested for the effect of collection site in the one species (*Micrurus lemniscatus*) that was collected from all localities, as there was otherwise high variability/stochasticity in which species were found at each collection site. If co-occurring species affect the anti-predator displays of individuals in a locality due to mimetic local adaptation, we would expect *M. lemniscatus* to have the highest behavioural variation because it was found sympatrically with every other species in our dataset. Finally, we tested for decay of the thrashing signal over the course of a trial by regressing thrash duration and sum of body curvature over time and assessing mean slope deviation from zero. All statistical models were built in R v 3.6.1 using the package ‘lme4’ (Bates et al. 2015) for mixed modelling, and significance was assessed at *α*=0.05 using number of groups in each GLMM as a conservative estimate of the degrees of freedom denominator.

To determine the thrashing frequency for each species, we divided the total number of observed thrashing events by the total amount of time each snake in the species was observed while encountering the experimental stimulus. The total amount of time excludes trials in which no thrashing was observed. Time in which the snake left the field of view was subtracted from this total time. To then compare these metrics across species, we bootstrapped data by randomly sampling with replacement from a combined dataset comprised of all individuals in a given species. To maintain the differences in propensity to thrash, these bootstrapped values were sampled in proportion to the thrashing frequency described above.

## Results

### Overall behavioural response

We recorded 160 behavioural trials in total across 25 snakes: 14 trials from 3 individuals of *Micrurus annellatus*, 16 trials from 2 individuals of *M. hemprichii*, 71 trials from 12 individuals of *M. lemniscatus*, 30 trials from 4 individuals of *M. obscurus*, and 29 trials from 4 individuals of *M. surinamensis* (Table 1).

Not all individuals responded to all predator cues in our trials. We found that the probability of response depended most strongly on an additive effect of both body size (binomial GLMM; *F*_*1,22*_= 5.732, *P*=0.026) and species (*F*_*4,22*_= 5.907, *P*=0.002), with larger individuals and *M. surinamensis* and *M. hemprichii* least likely to respond (Fig. 3). We did not find a significant effect of collection locality (*M. lemniscatus* only, see Methods; *F*_*2,12*_= 0.099, *P*=0.906) on the probability of response, nor significant interaction between SVL and species (*F*_*4,17*_= 0.665, *P*= 0.625). We also found no effect of sex (*F*_*1,22*_= 0.780, *P*= 0.387) or temperature (*F*_*1,25*_= 1.474, *P*= 0.236) on display probability, although we note that most of our trials were conducted across a narrow range of body temperatures between 25.1-27.8°C (q20-q80). Although our ability to run some higher order multiple regressions was limited by our sample sizes, the effect of stimulus type was marginally significant in a single fixed effect model (*F*_*2,22*_= 3.278, *P*=0.057) and may interact with other effects. Observationally, response to a contact stimulus produced a response in every individual tested, but the probability of response varied across species and body sizes in looming and vibration trials.

**Figure 3.**
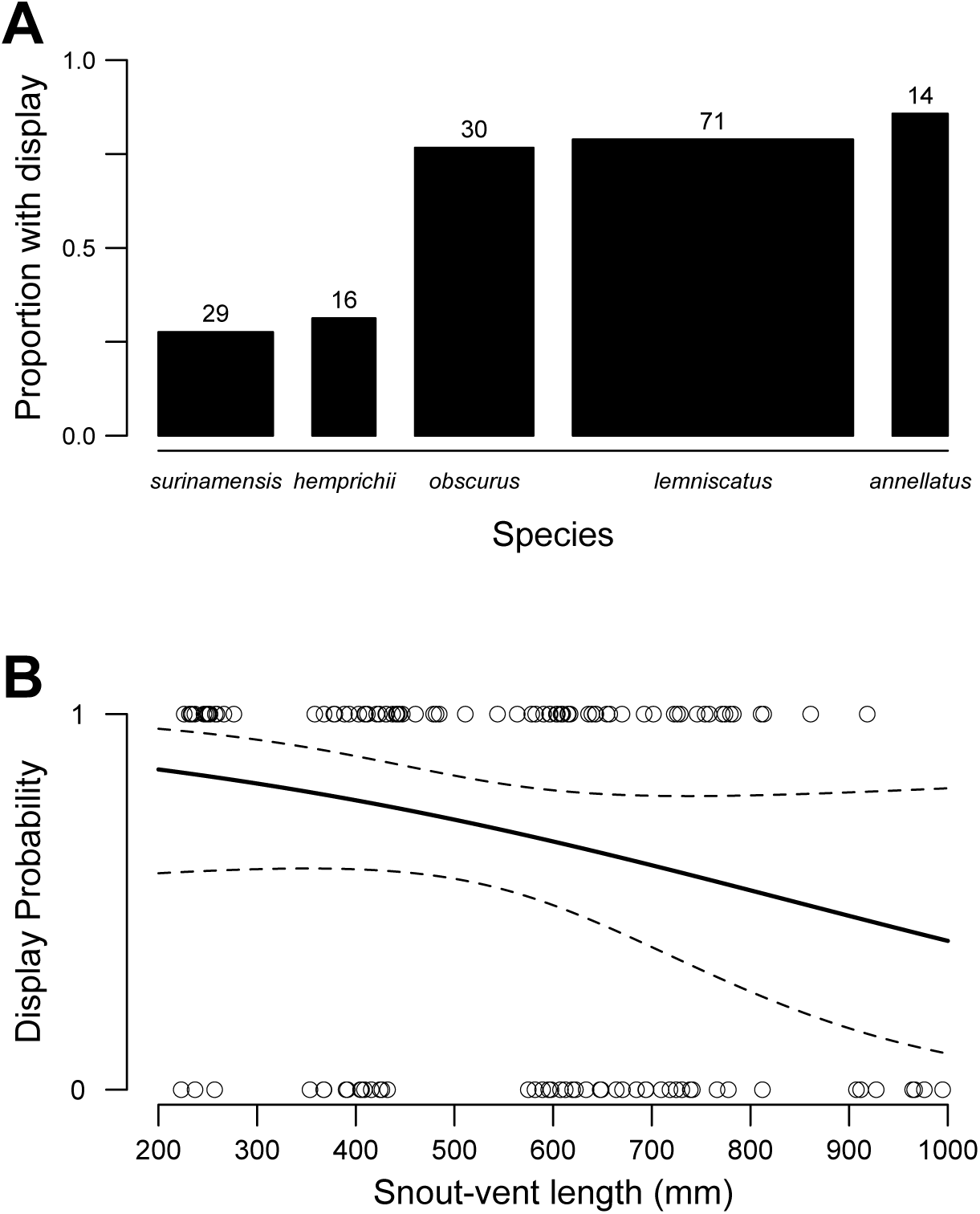
Not all individuals performed a thrashing display in response to all cues. Display probability depended most strongly upon (A) species identity (binomial GLMM: *F*_*4,22*_= 5.907, *P*=0.002) and (B) body size as measured by snout-vent length (SVL; *F*_*1,22*_= 5.732, *P*=0.026). Fitted effect of SVL is predicted from a model that includes collection ID as a random factor, as all individuals were tested more than once.

When analysing the thrashing display of the individuals that did respond to a predator cue, we found that thrash duration depended on body size (*F*_*1,20*_= 48.075, *P*<0.001) and stimulus type (*F*_*2,20*_= 4.877, *P*= 0.019), with larger snakes and those presented with contact stimuli performing longer thrash displays. We also found that magnitude of body curvature in the post-thrash pose depended upon stimulus (*F*_*2,20*_= 5.601, *P*= 0.012) and species (*F*_*4,20*_= 5.962, *P*= 0.003), such that *Micrurus lemniscatus* and those presented with a looming cue displayed the highest degree of body curvature (Supp. Fig. 1). Sex had no effect on either magnitude (*F*_*1,19*_= 0.212, *P*= 0.651) or duration (*F*_*1,19*_= 2.266, *P*= 0.149; one unvouchered, unsexed individual excluded). We also found no effect of temperature on magnitude (*F*_*1,20*_= 0.284, *P*= 0.600) or duration (*F*_*1,20*_= 0.155, *P*= 0.697). We found no significant preference within individuals for directionality (or “handedness”) in local maximum curvature, as nearly all individuals turned both heads and bodies in both directions during display (mean proportion of local maximum body curves to the right: 0.495; mean proportion of poses with right-kinked heads: 0.430, but 17/20 individuals had proportions between 0.2-0.8). We also found no significant effect of body size, species, or stimulus on the number of thrashing events within trials (all *P* > 0.05). Additionally, we saw no significant decay of the thrashing response over the course of trials, either in duration of thrash (mean slope: -0.003, range: -0.077 to 0.043) or degree of body curvature (mean slope: -0.288, range: -5.23 to 5.46). We did not have enough replicates to perform a robust test of locality effects.

### Intraspecific variation in M. lemniscatus

We compared the location and magnitude of body curvatures of the post-thrash pose among all individuals of each species. The variation among nine individuals of *M. lemniscatus* (Fig. 4) is emblematic of the intraspecific variation found in each of the other species as well (see Supp. Figs. 2-5, available online).

**Figure 4:**
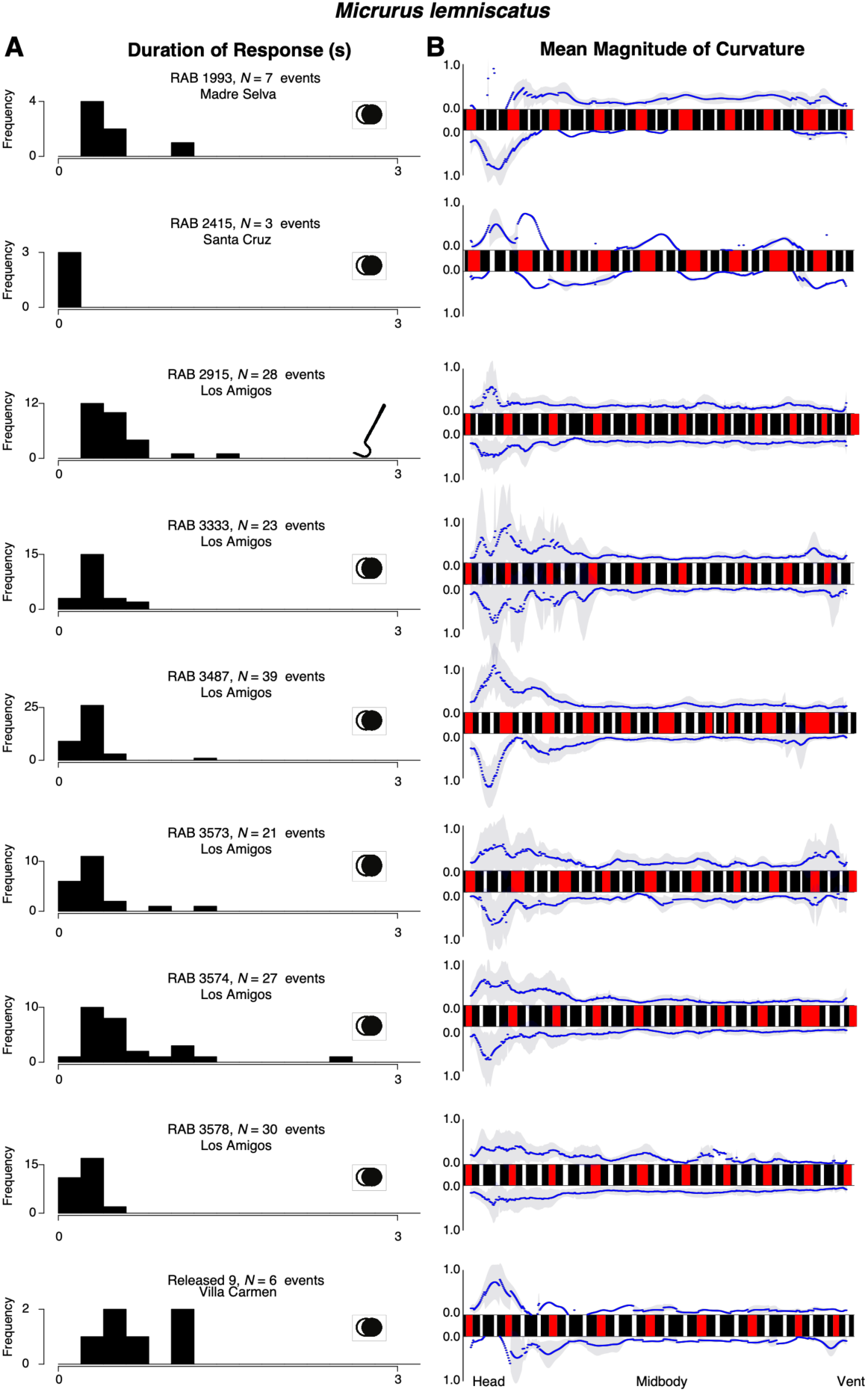
Intraspecific variation in thrash duration (A) and curvature (B) among individuals of *Micrurus lemniscatus*. For each row of plots, the histogram of thrash durations on the left and the mean magnitude of curvature plot on the right represent the same trial. The stimulus used in each trial is depicted by the icon on the right side of the histogram: two overlapping circles denote a looming stimulus and a snake hook denotes a tactile stimulus. Colour patterns of each snake reflect the observed colour patterns of each individual measured to sub-millimetre resolution. Snake patterns are arranged such that the anterior portion is towards the left.

Overall, there is strikingly high consistency in thrash duration (Fig. 4A) and location and degree of body curvature (Fig. 4B) both within and among individuals of this species. Areas of highest curvature (highest peaks) are concentrated towards the anterior portion in each snake, demonstrating the stereotypical “neck kinking” mentioned in qualitative descriptions of this display. In most individuals, both sides of the body are used fairly equivalently in body curving (Fig. 4B). However, within the bounds of this stereotyped display, the ability of snakes to dynamically adjust display components across cues and ontogeny contributed to substantial intraspecific variation across all metrics. The longest duration of thrash occurred in RAB 3574 (Fig. 4 A, seventh row), which was a generally active and moderately sized individual from Los Amigos. The most consistent and short durations of thrash were displayed by RAB 2415 (Fig. 4 A, second row), which was generally inactive, and the smallest snake captured of this species from Santa Cruz (overall SVL effect on thrash duration also shown in Supp. Fig. 1). As seen in the second row of Fig. 4B, this individual predominantly thrashed with the anterior portion of the body while keeping the rest of the body relatively stationary. There also appears to be substantial variation in how much of the anterior body displays the acute curves most associated with the neck, as well as how much of the posterior body displays the acute curves associated with tail coiling (not analysed here; see in particular RAB 3573 and 3333). Overall, there appears to be a similar amount of variation in behaviour as there is in coloration (precise colour variation shown in Fig. 4B, see legend), even though both traits are expected to be under exceptionally strong stabilizing selection.

### Interspecific variation

We found a similarly high consistency across species in the major features of the thrash display: the duration of thrashing tended to be relatively short (median value below 0.5 seconds for all species; Fig. 5A) and all species tended to have the largest magnitude of curvature towards the anterior portion of the body, irrespective of substantial variation in typical coloration and patterning (Fig. 5B; see also Supp. Figs 2-5). Beyond these major features, however, we again found variation in multiple components. The difference between the mean curvature at midbody and the mean curvature at the head was highest in *M. lemniscatus* and *M. obscurus*, and lowest in *M. hemprichii* and *M. surinamensis*. Similarly, the region of higher curvature extended posteriorly from the head much further in *M. lemniscatus* and *M. obscurus* than in *M. hemprichii, M. surinamensis*, and *M. annellatus*. However, posterior body curving near the tail was highest in *M. obscurus*, which happens to be the species with the shortest relative tail length in our dataset. This pattern may indicate compensation for a shorter tail by involving more of the body in the thrashing display. Additionally, the variance in thrashing durations was surprising, as longer bouts of thrashing were not equally likely for all species. *Micrurus annellatus* was the least likely to display for more than 1 second, *M. surinamensis* and *M. hemprichii* always displayed for less than 2 seconds, while *M. lemniscatus* and *M. obscurus* thrashing lasted up to 3 seconds (Fig. 5B).

**Figure 5:**
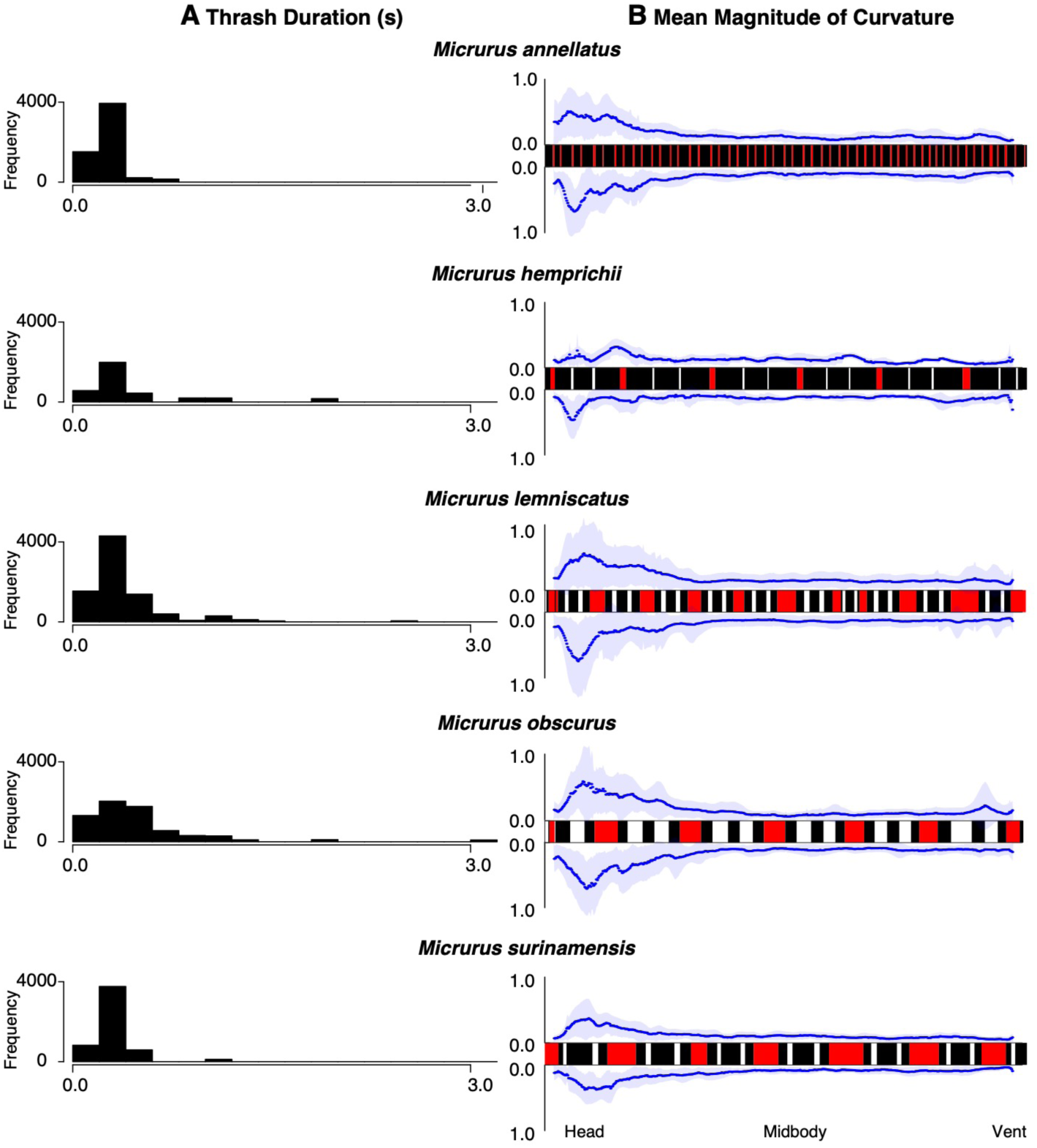
Interspecific variation in thrash duration (A) and curvature (B). Thrash durations for each species on the left are sampled with replacement proportionally by the ratio of events to total observed frames of video and plotted as histograms. On the right, the mean magnitude of curvature at each point along the body is plotted for one trial for each species in response to looming stimuli, with the exception of *Micrurus surinamensis*, which responded to a vibratory stimulus. Colour patterns of each snake reflect the observed colour patterns of each individual measured to sub-millimetre resolution. Snake patterns are arranged such that the left side is anterior.

## Discussion

By analysing the kinematics of non-locomotory coral snake anti-predator behaviours, we provide a new approach for quantitative comparison of critical signalling elements both within and among species. These results have important implications for testing theoretical expectations of mimicry systems and comparing results to other mimicry systems, such as butterflies, in which all players are chemically defended. Following theoretical predictions of these Müllerian systems, we expected to find greater behavioural similarity among sympatric species than intraspecific variation across multiple sites or even just high similarity across all individuals irrespective of species or locality. Contrary to this expectation, we found surprising diversity in 1) propensity to display at all, 2) duration of thrashing, and 3) dynamics of body positions and kinematics during the display, with no clear relationship to patterns of species sympatry. In particular, we found that this signal can be dynamically adjusted across ecological contexts, such that small snakes and those under physical contact by a predator are the most likely to respond and produce the most vigorous responses.

### Ecologically relevant variation in signal construction: what matters?

Our superficially paradoxical results add to mounting evidence that the drivers of trait variation in mimicry systems are not well understood (Joron and Mallet 1998; Mallet and Joron 1999; Cox and Davis Rabosky 2013; Davis Rabosky et al. 2016a). Although one interpretation could be that South American coral snakes are not part of a mimicry system, a more likely possibility is that the lethal levels of toxicity allow for greater trait variation due to the higher cost to predators that erroneously identify prey (Pough 1988). In particular, our results suggest that some aspects of the behavioural display – a short thrash duration and an acutely kinked neck – may be more important for effective signalling to predators than other traits, such as which direction the body curves or the degree of curvature beyond the neck. Additionally, our behavioural results provide an interesting comparison to variation in colour pattern, which is fairly extreme among species (Fig. 5B) but considered the most important trait for communicating the potential for chemical defence. More broadly, our ability to use a single method to analyse anti-predator display across a clade with a divergence date of at least 10 million years (Pyron and Burbrink 2014) gives some appreciation for how long the main thrashing element of this display has been maintained within the coral snake lineage. However, our results suggest that simplistic models of signal canalization in mimicry systems may benefit from expansion that accounts for the ways in which signals can and do still vary within and among species.

### Challenges of studying non-locomotory behaviour in venomous coral snakes

The non-locomotory movements of snakes present several unique challenges to quantitative biomechanical analyses. While many forms of snake locomotion involve either linear or roughly sinusoidal body shapes (Lissmann 1950; Jayne 1986; Moon and Gans 1998; Hu et al. 2009; Socha 2011, Titcomb et al. 2013), non-locomotory behaviours are less mechanically constrained and therefore result in a high frequency of self-occlusion. The elongate body form and high degrees of freedom conferred by the highly articulated musculoskeletal system result in more acute curves than observed in other model organisms with elongate body forms (e.g. *Caenorhabditis elegans*, Padmanabhan 2012; Brown 2013). Such extreme self-occlusions make it difficult to use automated tools for passively tracing the snake centreline.

Furthermore, the banded colour patterns and self-mimicry by the tail of the head (Greene 1973) make it difficult to identify the morphological features of a snake from an isolated video frame without context. These features likely contribute to snake fitness by making it difficult for a predator to target an attack towards the head (Brodie III 1993) and have resulted in a particularly challenging dataset for existing computer vision tools.

While fiduciary markers placed on the snake can aid in collecting precise kinematic data (Gart 2019), the handling required to affix such markers likely alters the behavioural response of the snake. Furthermore, the highly toxic venom of these snakes, together with impressively adhesive-resistant skin lipids (Torri et al. 2014), makes it impractical and unsafe to attempt such marking.

The smooth surface of the arena required for thorough cleansing resulted in lower substrate friction for the snake. Although some slipping was observed, recent studies suggest that snake motor control is independent of surface friction (Schiebel et al. 2019). Thus, we argue that our results likely reflect snake behavioural variation on more natural substrates.

### Future directions

We demonstrate that a video-based approach can be feasibly applied to quantitatively examine the non-locomotory behaviours of snakes under semi-natural conditions. This approach facilitates ecologically relevant biomechanical inquiry with strong evolutionary impact. While we recognize that our stimuli, especially the vibratory stimulus, may not be a perfect match to those provided by potential predators, at least one individual responded to every category of stimulus with an anti-predator display.

In addition to venomous coral snakes, distantly related snakes with varying toxicity have independently converged several times on these conspicuous colour patterns and thrashing displays (Greene 1979; Davis Rabosky et al. 2016b). Just as previous studies have leveraged a quantitative analysis of colour pattern to measure convergence and experimentally examine the effect on predation rates (Brodie III 1993; Buasso 2006), the analysis presented above enables more precise measurement of convergence and an experimental approach for determining the effect of behaviour on predation rates. Since snakes of all sizes, and consequently all ages, display vigorous thrashing behaviour in the absence of any parental care (Shine 1988), this anti-predator response has likely evolved as an innate response, with little learning over the course of a lifetime. In this case, quantitative characterization of anti-predator behaviours can be modelled as phenotypic traits to gain insight into the evolutionary processes underlying the patterns of behaviour observed in nature.

## Author contributions

TYM, JGL, and ARDR designed the experiment and collected the data. TYM, ARDR, and SMD analysed the data. TYM and ARDR wrote the main draft of the manuscript, and all authors contributed to manuscript editing and revision.

## Acknowledgements

The authors thank the many members of the University of Michigan/Peru Herpetology Expeditions for their assistance with capturing the snakes. Funding for field collection expeditions was provided by the University of Michigan to TYM and ARDR and a Packard Foundation Fellowship to Dan Rabosky. Funding for collaboration and analysis was provided by a University of Michigan MCubed grant. We particularly thank Ciara Sanchez Paredes, Briana Sealey, and Erin Westeen for leading several seasons of behavioural filming in the field, and Rudi von May for collection and export permit acquisition. Peter Cerda, John David Curlis, Courtney Whitcher, Daniel Nondorf, and Molly Hirst were especially helpful labelling still frames from videos. We also thank Ashley Thompson and Timothy Renney for assistance with illustrations. The authors declare no conflicts of interest.

## Data accessibility

Videos will be archived and freely available at https://deepblue.lib.umich.edu/data. Code to compute the curvature of the snakes is posted in a GitHub repository.

## Notes

https://doi.org/10.7302/bksb-t580

