## Supporting Information for "A kinematic analysis of *Micrurus* coral snakes reveals unexpected variation in stereotyped anti-predator displays within a mimicry system"

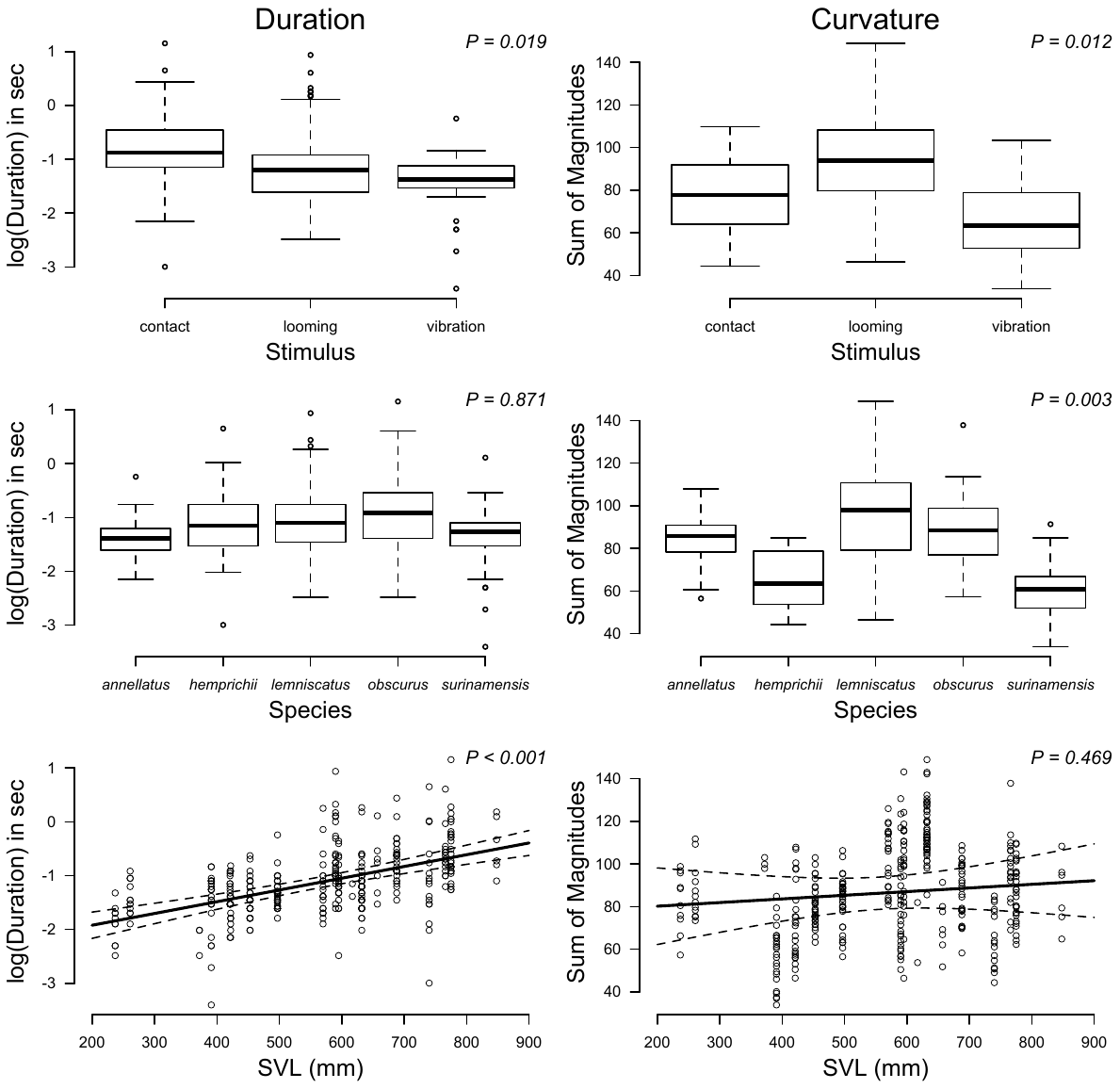

**Supporting Figure 1:** Different factors affected different aspects of coral snake thrashing displays. Linear mixed effects models (LMM) that included individual collection ID as a random effect showed that the type of stimulus had significant effects on thrashing, with contact evoking the longest thrash durations (*F_2,20_*= 4.877, *P*=0.019) and looming evoking the curviest post-thrash postures (*F_2,20_*= 5.601, *P*=0.012). Duration of thrashing bouts was not affected by species (*F_4,20_*= 0.306, *P*=0.871) but was greatly affected by the size of the snake (*F_1,20_*= 48.075, *P*<0.001). Curvature was highly subject to species identity (highest in *M. lemniscatus* and lowest in *M. hemprichii* and *M. surinamensis*; *F_4,20_*= 5.962, *P*=0.003) but was not dependent on the size of the snake (*F_1,20_*= 0.545, *P*=0.469). Sex had no effect on either magnitude (*F_1,19_*= 0.212, *P*= 0.651) or duration (*F_1,19_*= 2.266, *P*= 0.149; one unvouchered, unsexed individual excluded).

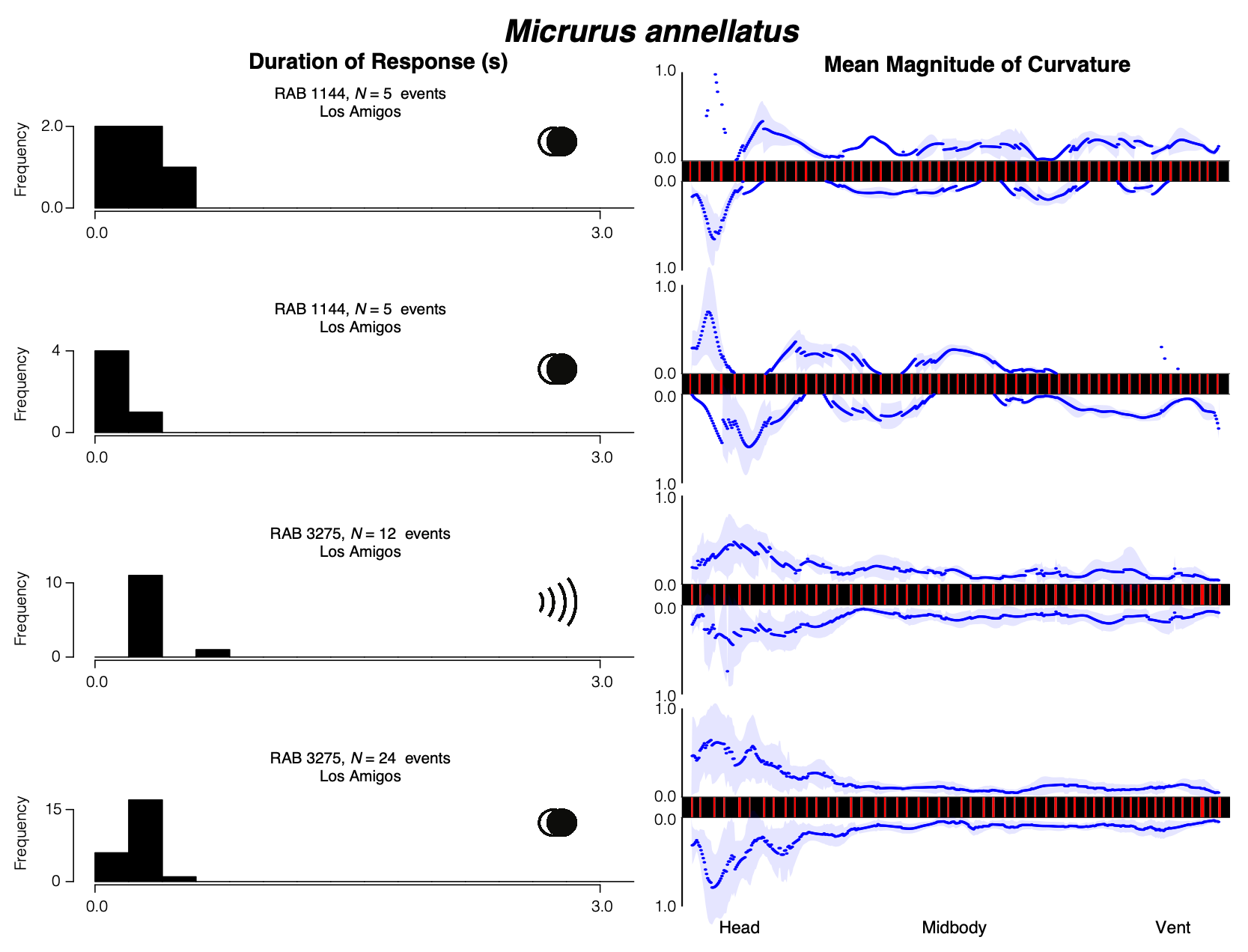
**Supporting Figure 2:** All trials from all individuals of *Micrurus annellatus* that included thrashing. Histograms of thrash duration are shown on the left and the mean magnitude of curvature at every point along the body is shown on the right. The stimulus used in each trial is depicted by the icon on the right side of the histogram: two overlapping circles denote a looming stimulus and a wave transmission icon denotes a vibration stimulus. Colour patterns of each snake reflect the observed colour patterns of each individual measured to sub-millimetre resolution. Snake patterns are arranged such that the anterior portion is towards the left.

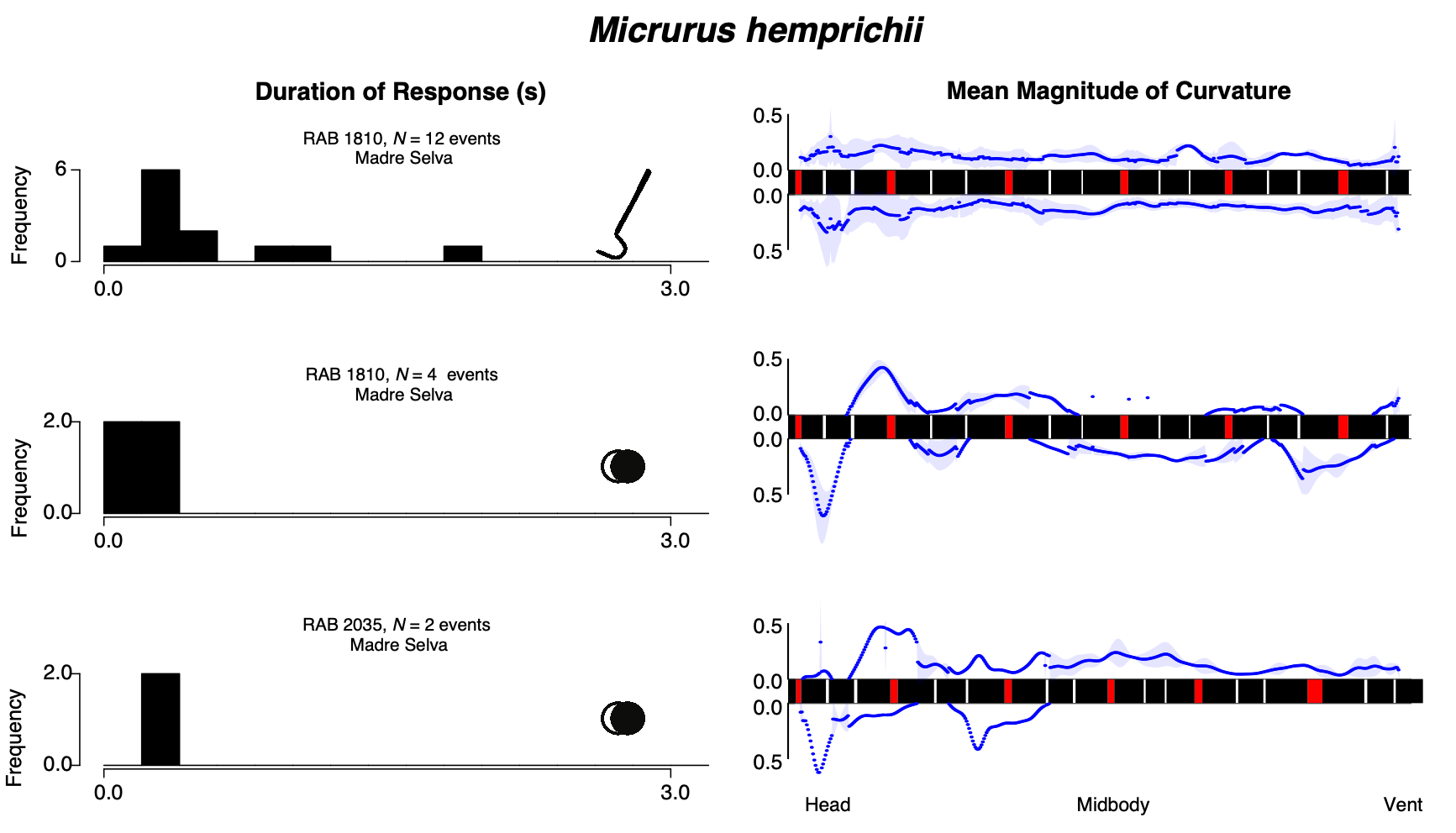
**Supporting Figure 3:** All trials from all individuals of *Micrurus hemprichii* that included thrashing. Histograms of thrash duration are shown on the left and the mean magnitude of curvature at every point along the body is shown on the right. The stimulus used in each trial is depicted by the icon on the right side of the histogram: two overlapping circles denote a looming stimulus and a snake hook denotes a tactile stimulus. Colour patterns of each snake reflect the observed colour patterns of each individual measured to sub-millimetre resolution. Snake patterns are arranged such that the anterior portion is towards the left.

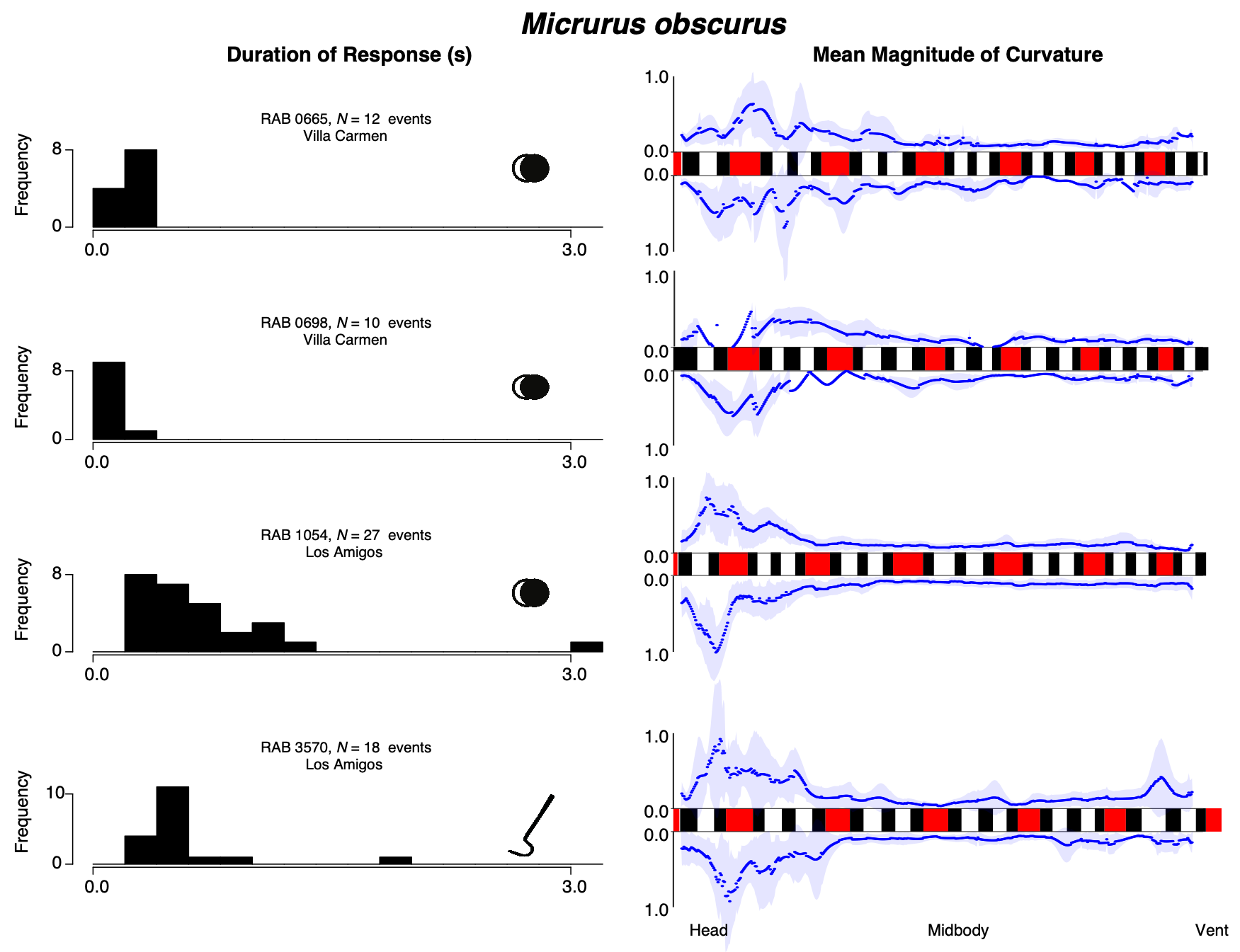
**Supporting Figure 4:** All trials from all individuals of *Micrurus obscurus* that included thrashing. Histograms of thrash duration are shown on the left and the mean magnitude of curvature at every point along the body is shown on the right. The stimulus used in each trial is depicted by the icon on the right side of the histogram: two overlapping circles denote a looming stimulus and a snake hook denotes a tactile stimulus. Colour patterns of each snake reflect the observed colour patterns of each individual measured to sub-millimetre resolution. Snake patterns are arranged such that the anterior portion is towards the left.

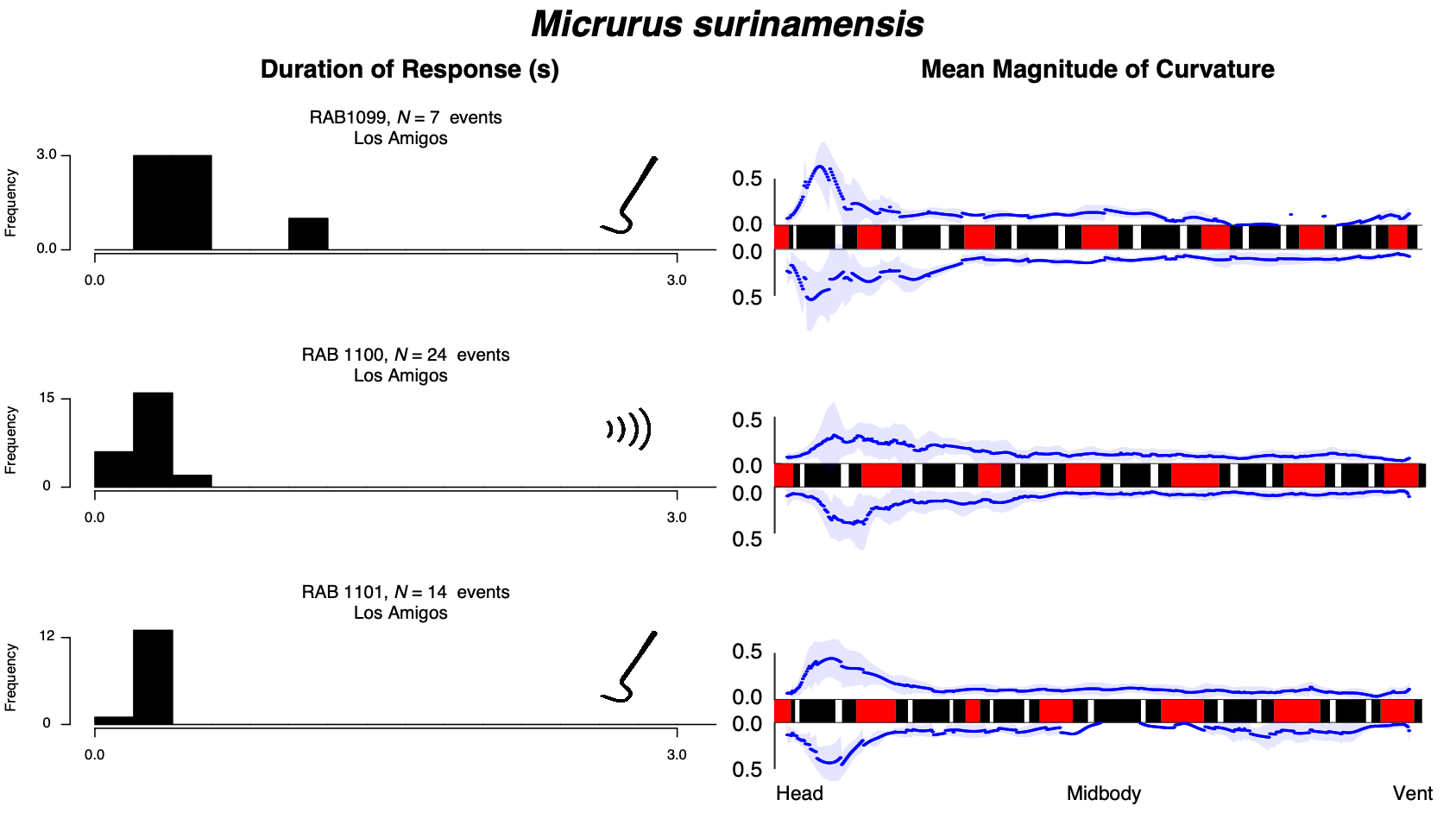
**Supporting Figure 5:** All trials from all individuals of *Micrurus surinamensis* that included thrashing. Histograms of thrash duration are shown on the left and the mean magnitude of curvature at every point along the body is shown on the right. The stimulus used in each trial is depicted by the icon on the right side of the histogram: a snake hook denotes a tactile stimulus and a wave transmission icon denotes a vibration stimulus. Colour patterns of each snake reflect the observed colour patterns of each individual measured to sub-millimetre resolution. Snake patterns are arranged such that the anterior portion is towards the left.

**Supporting Video 1:** Video of thrashing anti-predator behavior exhibited by an adult female *Micrurus lemniscatus* (RAB 3487) from Los Amigos Biological Station in southern Peru in response to a looming stimulus produced by a white cloth. This video has been undistorted and rectified. Note the frequent self-occlusions, the intermittent nature of the thrashes, and the similarity in shape between the kinked head and coiled tail.

[Provided as a separate file]
